# Anatomical Region-Specific Transcriptomic Signatures and the Role of Epithelial Cells in Pterygium Inflammation: A Multi-Omics Analysis

**DOI:** 10.1101/2024.11.25.625095

**Authors:** Chaerim Song, Seokho Myung, Hanseul Cho, Tae Gi Kim, Soohyun Chun, Minju Seo, Hyunmin Yu, Seoyeon Kim, Ye-Ah Kim, Junghyun Kim, Jaeyong Shin, Jaheon Kang, Yoonsung Lee, Min Seok Kang, Man S Kim

## Abstract

**Background:** The anatomy of pterygium consists of the head, neck, and body. Prior studies have examined the three parts collectively, yet the treatment methods and their success rates differ based on the anatomical characteristics. In this study, we divide the pterygium tissue into Main (head and neck) and Acc (body) to investigate its pathogenesis across regions.

**Purposes:** This study aims to understand the pathogenesis and identify potential therapeutic targets of pterygium. Through a robust multi-omics analysis of bulk and single-cell RNA sequencing (RNA-seq) data from pterygium patients, we focus on the expression of inflammatory and mitochondrial energy pathways and identify potential genes responsible for the upregulated pathways in pterygium.

**Methods:** We collected bulk RNA-seq data from six pterygium patients and single-cell RNA-seq data from two pterygium patients. We then investigated the pathway enrichment, pathway correlation, differential gene expressions, protein-protein interactions, and cell-cell communications of pterygium.

**Results:** From the analysis of bulk RNA-seq data, the distribution of sample points from the principal component analysis plot and the pathways enriched from the Gene Ontology analysis showed distinct expression patterns in the Acc group compared to the Main and control groups. This suggested the need to separate the Main and Acc regions within pterygium samples and utilize single-cell RNA-seq data to understand the differences between the Main and control groups that the bulk data could not capture. The annotation of integrated single-cell data revealed a cluster of epithelial cells containing only pterygium samples. Cells from this cluster exhibited significant contributions to the ANGPTL, IL1, and KLK signaling networks in the cell-cell communication analysis. We also observed significant upregulation of the cells in the inflammatory pathways related to integrated stress response and the renin-angiotensin-aldosterone system, both of which showed high correlations with energy metabolism pathways. Significant changes in the expression of multiple pro-inflammatory, antioxidant, and immune-related genes were also identified.

**Conclusion:** The different expression patterns between the Main and Acc groups suggest the need to consider different anatomical regions separately in future studies of pterygium. Additionally, the significant role epithelial cells from the Main group play in the inflammation of pterygium presents a potential clinical approach to the disease.

## 1. INTRODUCTION

Pterygium is characterized by a triangular, wedge-shaped tissue growth extending into the cornea. One primary contributing factor is exposure to ultraviolet (UV) radiation. Histopathologically, it is marked by elastotic degeneration of the conjunctival collagen [1]. Many studies have recently been conducted to determine the exact causes and mechanisms of pterygium. Previously, we identified that the intrinsic gene regulatory mechanisms are closely linked to the inflammatory reactions of pterygium using RNA sequencing, and compared regulatory patterns of functional genes or gene modules of pterygium between Asian and European ethnic groups [2].

The main treatment of pterygium is surgical removal. However, recurrence remains the main obstacle to effective treatment, characterized by the regrowth of fibrovascular tissue across the limbus into the cornea. Various surgical techniques have been employed, but none are universally favored due to inconsistent recurrence rates. Regardless of the chosen method, removing the pterygium is the initial step in the procedure. Many ophthalmologists prefer to detach the pterygium head from the underlying cornea [3].

The anatomical characteristics of pterygium affect the treatment method and its success rate. The anatomy of pterygium can be divided into three parts: apex or head, neck, and body [4]. The invading portion growing toward the center of the cornea which contains the apex of the tissue is called the head. The head is a vascular area firmly attached to the cornea and can affect the corneal epithelium and the Bowman’s layer, potentially leading to vision impairment. In particular, the leading edge of the head is called the cap. The communicating part between the body and the head, which overlies the limbus, is named the neck. The conjunctival portion with a base toward the medial canthus is known as the body. The likelihood of recurrence is high if the head of the pterygium is not completely removed during surgery, especially when the pterygium has invaded the Bowman’s layer. Procedures such as autologous conjunctival grafting or amniotic membrane transplantation are also used to prevent recurrence. If the body of pterygium is not adequately managed, the risk of recurrence increases [5]. These points therefore suggest that the pathogenesis or inflammatory responses of pterygium may differ based on its anatomical regions.

This study aims to understand the pathogenesis and identify potential therapeutic targets of pterygium through the analysis of RNA sequencing data. Specifically, we seek to investigate the differences between the anatomical parts of the pterygium and explore which factors may influence the onset and recurrence of pterygium.

## 2. RESULTS

Since no prior study has examined the gene expression profiles of pterygium across different anatomical regions, we categorized the pterygium samples into two groups: the Main group (Cap, Head, Neck) and the Acc group (Body). We first identified significantly differentially expressed genes (DEGs) among the Main, Acc, and Normal groups. A relatively higher number of DEGs (25468 in Main vs. Acc; 24785 in Normal vs. Acc) were found when the Main and Normal groups were analyzed against the Acc group, compared to the 21887 DEGs found in Main vs. Normal (Fig. 1a). The principal component analysis (PCA) plot also revealed a clear separation of sample points between Acc and the other two groups, suggesting the presence of distinct genetic expression patterns between the Acc and Main regions (Fig. 1b). This finding points out the need for the separation of pterygium regions, unlike previous studies that analyzed the Acc and Main regions as a single group [6].

**Figure 1.**
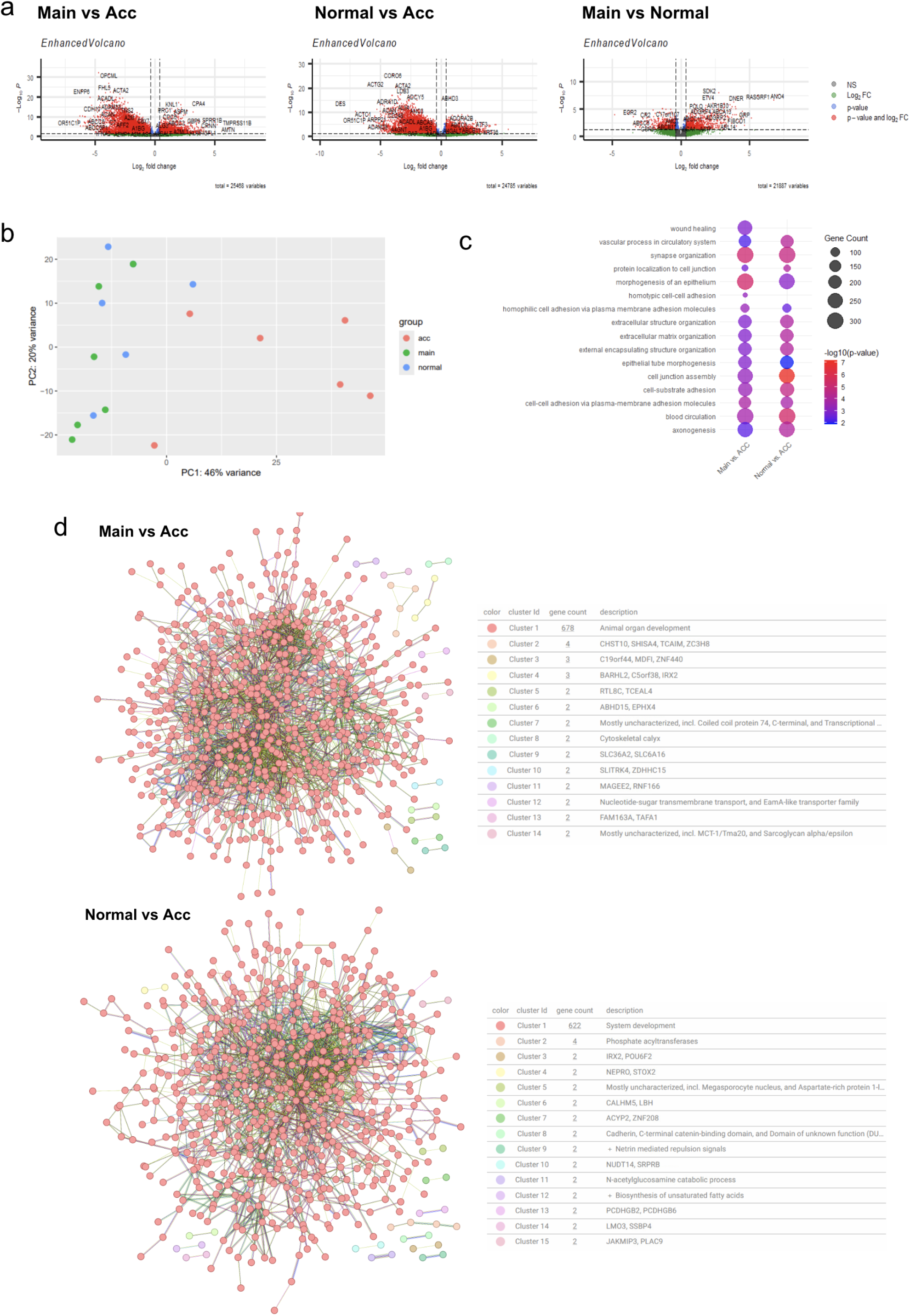
**(a)** Volcano plots of DEGs from pairwise comparisons among the Main, Normal, and Acc groups (p-value < 0.05, log2 fold-change > 0.38). **(b)** PCA plot of PC1 and PC2 showing separation of Acc group from the Main and Normal groups. **(c)** Most commonly enriched pathways from GO analysis. **(d)** PPI networks with top 1000 significant DEGs.

To assess whether the identified DEGs are linked to functions documented in previous studies, we conducted a Gene Ontology (GO) analysis (Fig. 1c). Pathways including synapse organization, epithelium morphogenesis, and cell junction assembly, commonly known to be related to organ and system development, were significantly enriched in both the Main and Normal groups relative to the Acc group.

To explore the functional roles of DEGs enriched in the Main and Normal groups compared to the Acc group at the proteomic level, we analyzed protein-protein interaction (PPI) networks using STRINGdb. As shown in Figure 1d, a prominent cluster associated with organ development emerged in the comparison of Main and Acc, encompassing pathways like collagen fibril organization, regulation of vascular permeability, and extracellular matrix organization. The PPI in the comparison between Normal and Acc also displays a major cluster associated with system development, including pathways such as muscle cell differentiation, and vasculogenesis. These observations are in line with the findings from the GO analysis, highlighting the distinction between the Acc and Main group, and relatively similar characteristics between the Main and Normal groups.

Inflammation and oxidative stress are key factors in the pathology of pterygium [2]. To further investigate the association between the inflammatory pathways and anatomical region, the normalized enrichment score (NES) was calculated for each pathway using Fast Gene Enrichment Analysis (fGSEA) with a customized inflammatory gene list obtained from [7]. As shown in Figure 2a, most inflammatory modules were generally upregulated in the Main and Normal groups compared to the Acc group. Notably, three modules—Interleukins in Adaptive Immunity, Complement Activation/Fibrin Deposition, and the AGT Regulator Axis in the Renin-Angiotensin-Aldosterone System (RAAS)—were downregulated in both the Main and Normal groups.

**Figure 2.**
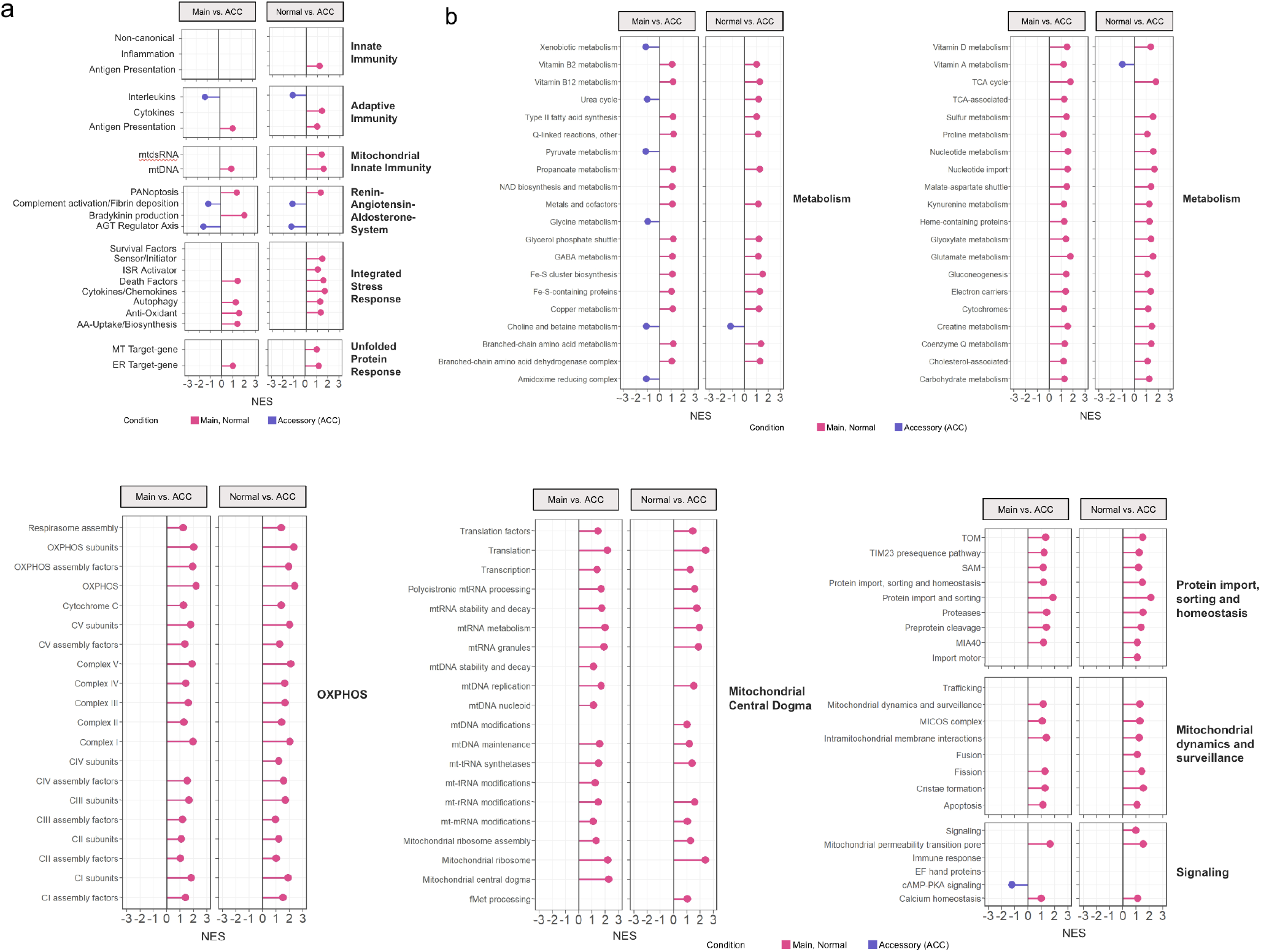
Lollipop plots comparing the Acc group with the Main and Normal groups. Generated with NESs computed with core inflammatory and mitochondrial genes. **(a)** Lollipop plot of inflammatory pathways. **(b)** Lollipop plots of mitochondrial energy pathways.

Given the critical role of mitochondria in inflammation and oxidative stress, we further examined the expression profiles of core genes in energy-related mitochondrial pathways in the Acc group using MitoCarta3.0 (Fig. 2b) [8, 9]. Consistent with the inflammatory profile, most mitochondrial pathways, including oxidative phosphorylation (OXPHOS), mitochondrial central dogma (MCD), mitochondrial dynamics and surveillance, signaling, protein import, and homeostasis, were upregulated in the Main and Normal groups compared to the Acc group. The majority of metabolism-related pathways were also upregulated in the two groups except for the pathways involved in pyruvate metabolism, xenobiotic metabolism, the urea cycle, glycine metabolism, choline and betaine metabolism, and the amidoxime reducing complex. The clear distinction between the Acc and Main groups and the relatively similar up- and down-regulation patterns between the Main and Normal groups support our point that the anatomical region of the samples is highly related to its transcriptional profile. This therefore suggests the need to separate Acc and Main groups in future pterygium studies.

In order to provide an overview of the molecular and cellular landscape of patient-matched selective pterygium samples and Normal samples, we performed single-cell RNA sequencing (scRNA-seq) profiling with 10X Chromium from freshly resected pterygium samples and their adjacent control (cornea) samples from two patients. Due to the transcriptional differences we identified between the anatomical regions of pterygium, we selectively collected the Main samples (non-ACC) to represent pterygium. The samples were integrated using joint analysis of heterogeneous samples (Fig. 3a). Using marker genes from Zhang et al., we identified 7 distinct clusters of cell types, including epithelial cells (KRT7, KRT5, KRT19), endothelial cells (RAMP3, VWF, PECAM1), fibroblasts (COL1A1, COL1A2, DCN), T/NK cells (TRAC, CD3E, CD3D), myeloid cells (CD68, LYZ, TYROBP), and melanocytes (TYRP1, PMEL, MLANA) [10]. Interestingly, we identified a cluster that only consisted of pterygium samples among the cells annotated as epithelial cells and denoted the group separately as pterygium-specific epithelial cells. This indicates that this cell type may be unique to or highly associated with the Main group and likely plays a role in the pathology of pterygium.

**Figure 3.**
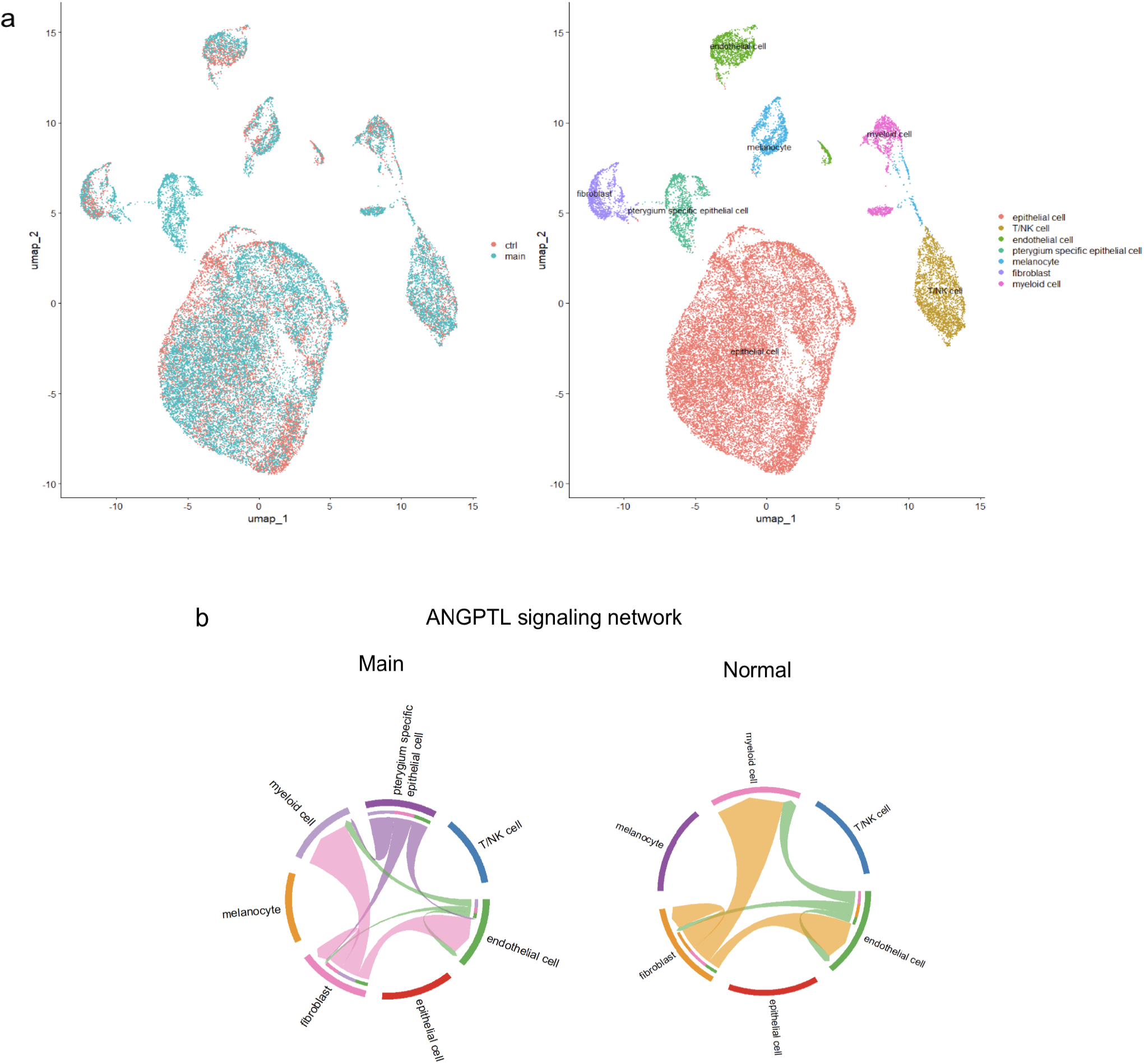
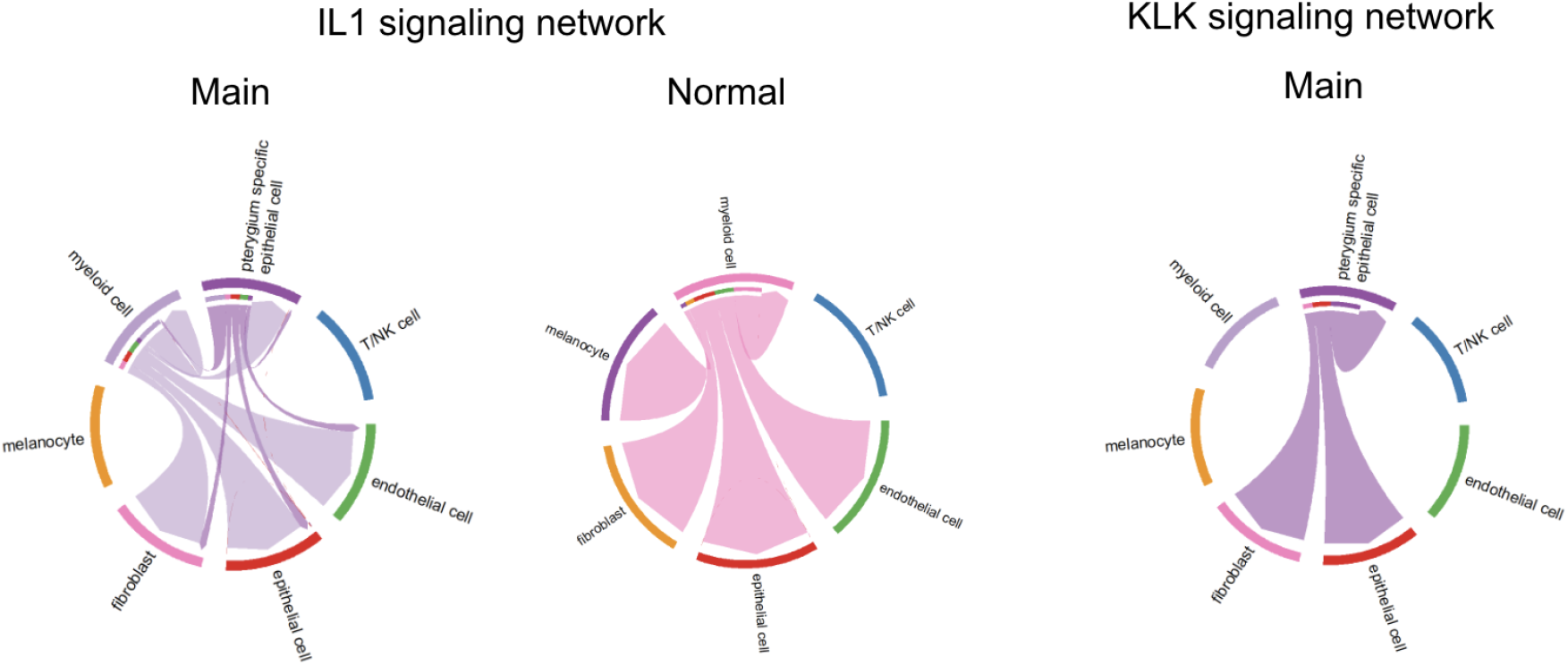
**(a)** Clustering of integrated data (left) and clusters annotated by their cell types (right). **(b)** Cell-cell communication of inflammatory signaling networks that showed significant contributions of pterygium-specific epithelial cells in the interactions between cell types.

To examine the differences in how the Main and Normal samples communicate at a cellular level, we analyzed the interactions between different cell types within various inflammation-related signaling networks using CellChat. As illustrated in Figure 3b, pterygium-specific epithelial cells significantly contributed to the inflammatory response in pterygium through the ANGPTL, IL1, and KLK signaling networks. Fibroblasts, myeloid cells, endothelial cells, and epithelial cells showed interactions with pterygium-specific epithelial cells in these pathways (Fig. 3b).

To estimate the activation levels of inflammatory pathways in each cell type, we conducted fGSEA and obtained the corresponding NESs using a customized inflammatory gene list obtained from [7] (Fig. 4a). Compared to the cells in the control samples, the epithelial cells in pterygium samples were up-regulated in surface marker/receptor signaling in adaptive immunity and inflammation in innate immunity, whereas the canonical pathways in innate immunity were upregulated in both the epithelial and pterygium-specific epithelial cells. Unlike the epithelial cells found in both the pterygium and control samples, pterygium-specific epithelial cells were significantly upregulated in pathways involved in antioxidant responses and AA-uptake/biosynthesis from the Integrated Stress Response (ISR) as well as in complement activation/fibrin deposition and bradykinin production from the renin-angiotensin-aldosterone system (RAAS) (Fig. 4a). RAAS is known to be associated with retinal vasculopathy and inflammation [11], suggesting that it may be one of the pathways that the pterygium-specific epithelial cells contribute to in the pathology of pterygium.

**Figure 4.**
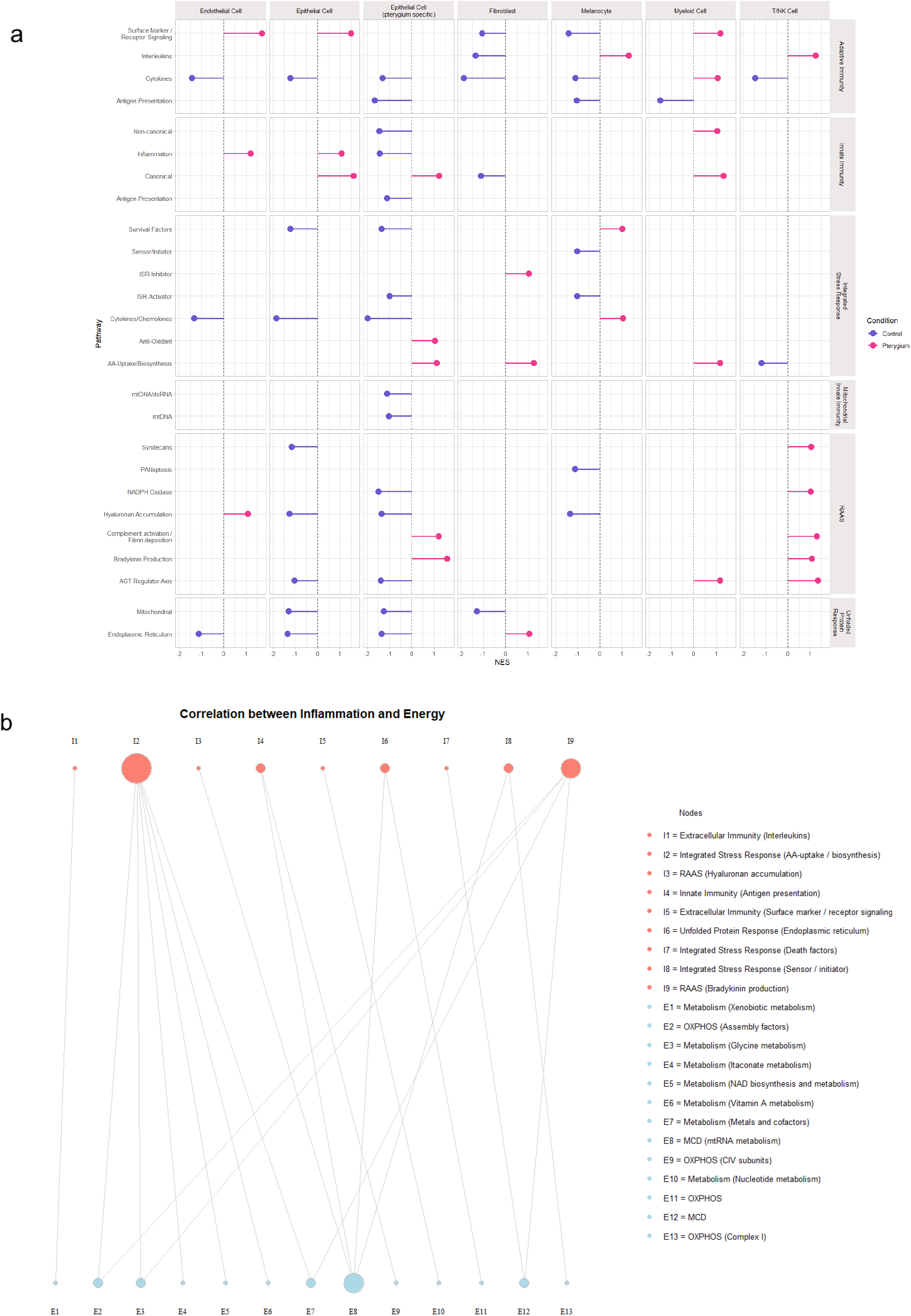
**(a)** Lollipop plot showing NES values (|NES| > 1) of inflammatory pathways for each cell type. **(b)** Correlation between inflammatory (pink) and mitochondrial energy (blue) pathways.

In T/NK cells, interleukin activity in adaptive immunity and most of the RAAS system pathways were upregulated in the pterygium samples. Endothelial cells exhibited upregulation of surface marker/receptor signaling in adaptive immunity. Fibroblasts showed increased ISR inhibitor and AA-uptake/biosynthesis in the ISR system. Interleukin activity in adaptive immunity as well as survival factors and cytokines/chemokines in the ISR were activated in melanocytes. In myeloid cells, surface markers and cytokines in adaptive immunity, both non-canonical and canonical pathways in innate immunity, AA-uptake/biosynthesis in the ISR, and the AGT regulator axis in the RAAS system were all upregulated (Fig. 4a).

To investigate the energy sources driving the upregulated inflammatory pathways, we analyzed the correlation between inflammatory and mitochondrial energy pathways using normalized enrichment scores (NES) from upstream analyses (Fig. 4b). Due to the skewed data distribution observed in the lollipop plot (Fig. 4a), a non-parametric method, the Van der Waerden test, was employed. Inflammatory pathways that exhibited significant correlations with the mitochondrial energy pathways include interleukins and surface marker receptor signaling in extracellular immunity, hyaluronan accumulation of RAAS, antigen presentation involved in innate immunity, endoplasmic reticulum pathways of unfolded protein response, and death factors and sensor/initiator pathways of ISR. Likewise, the mitochondrial energy pathways that showed significant correlations with the inflammatory pathways are metabolism pathways (xenobiotic metabolism, glycine metabolism, itaconate metabolism, NAD biosynthesis and metabolism, vitamin A metabolism, metals and cofactors, and nucleotide metabolism), mtRNA metabolism in the MCD, and CIV subunits and Complex I pathways of OXPHOS.

The inflammatory pathways that showed the highest correlation were AA-uptake/biosynthesis in ISR connected to six energy nodes, followed by bradykinin production in RAAS connected to five energy nodes. These two pathways were both upregulated in the pterygium-specific epithelial cells, suggesting that these cells are potentially related to the inflammatory responses highly correlated with the mitochondrial metabolism of pterygium tissues. Fibroblasts and myeloid cells were also upregulated in AA-uptake/biosynthesis, while T/NK cells were upregulated in bradykinin production (Fig. 4a).

To estimate the activation levels of mitochondrial energy pathways in each cell type, we also conducted fGSEA and obtained the corresponding NESs using MitoCarta3.0 (Fig. 5). We focused our analysis on metabolism, OXPHOS, and MCD, as these three showed the highest correlations with inflammatory pathways from Figure 4a. In general, the pterygium samples were downregulated in OXPHOS and MCD pathways compared to the control samples. The metabolism pathways, on the other hand, were more upregulated, suggesting that the metabolic reprogramming towards metabolic pathways in pterygium. This pattern corresponds to the previous study that reported OXPHOS to be upregulated in European pterygium samples while being downregulated in Asian samples [2].

**Figure 5.**
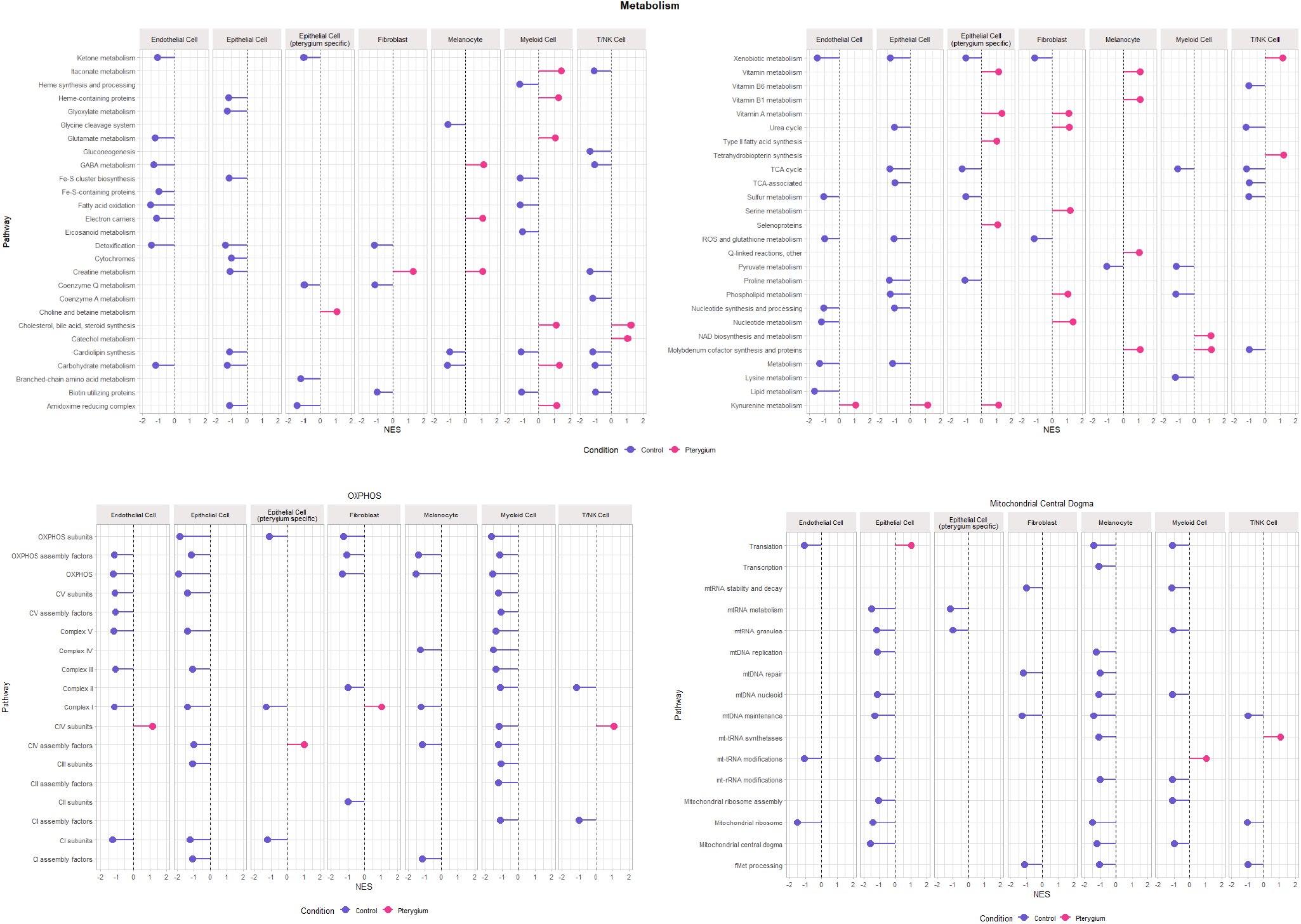
Lollipop plots showing NES values (|NES| > 1) of metabolism, OXPHOS, and MCD pathways of each cell type.

Metabolic pathways that were upregulated in the pterygium-specific epithelial cells include choline and beta metabolism, vitamin metabolism, vitamin A metabolism, type II fatty acid synthesis, selenoproteins, and kynurenine metabolism. Fibroblasts of pterygium samples showed upregulations in creatine metabolism, vitamin A metabolism, urea cycle, serine metabolism, phospholipid metabolism, and nucleotide metabolism. Melanocytes were upregulated in GABA metabolism, electron-carriers, creatine metabolism, vitamin metabolism, vitamin B1 metabolism, Q-linked reactions, and molybdenum cofactor synthesis and proteins. The pathways upregulated in the T/NK cells of pterygium samples include cholesterol, bile acid, steroid synthesis, catechol metabolism, xenobiotic metabolism, and tetrahydrobiopterin synthesis. Endothelial cells and epithelial cells (excluding pterygium-specific epithelial cells) were mostly downregulated in the metabolic pathways except for kynurenine metabolism.

Among the OXPHOS pathways, the Complex IV subunits in the endothelial cells. Complex IV assembly factors in the pterygium-specific epithelial cells, Complex I in fibroblasts, and Complex IV subunits in T/NK cells were upregulated in the pterygium samples relative to the control group. Among the MCD pathways, translation in epithelial cells, mt-tRNA modification in myeloid cells, and mt-tRNA synthetases in T/NK cells were upregulated.

Having found a crucial association between pterygium-specific epithelial cells and core inflammatory and mitochondrial energy pathways, we also examined the expressions of the specific genes involved. As shown in Figures 6a and 6b, pterygium-specific epithelial cells showed the most striking alterations. The significantly altered genes related to inflammation that were upregulated in pterygium-specific epithelial cells include: (i) STAT1 and DDX60, which act in sustained pro-inflammatory responses [12, 13]; (ii) FAS, which actively remove damaged cells, a process likely driven by chronic inflammation [14]; (iii) SLC7A11 and NQO1, related to antioxidant defenses in response to chronic oxidative stress [15, 16]; (iv) BDKRB1 and BDKRB2, which encode bradykinin receptors and are involved in the inflammatory environment; (v) THBD (thrombomodulin) and SDC1 (syndecan-1), involved in ongoing tissue repair and remodeling [17]. The genes that were downregulated include: (i) CX3CL1, CSF3, IL6, and CXCL8, which suggest an overall impairment in immune cell recruitment [18, 19]; (ii) PPARGC1A and SOD2, which support mitochondrial dysfunction [20] (Fig. 6a).

**Figure 6.**
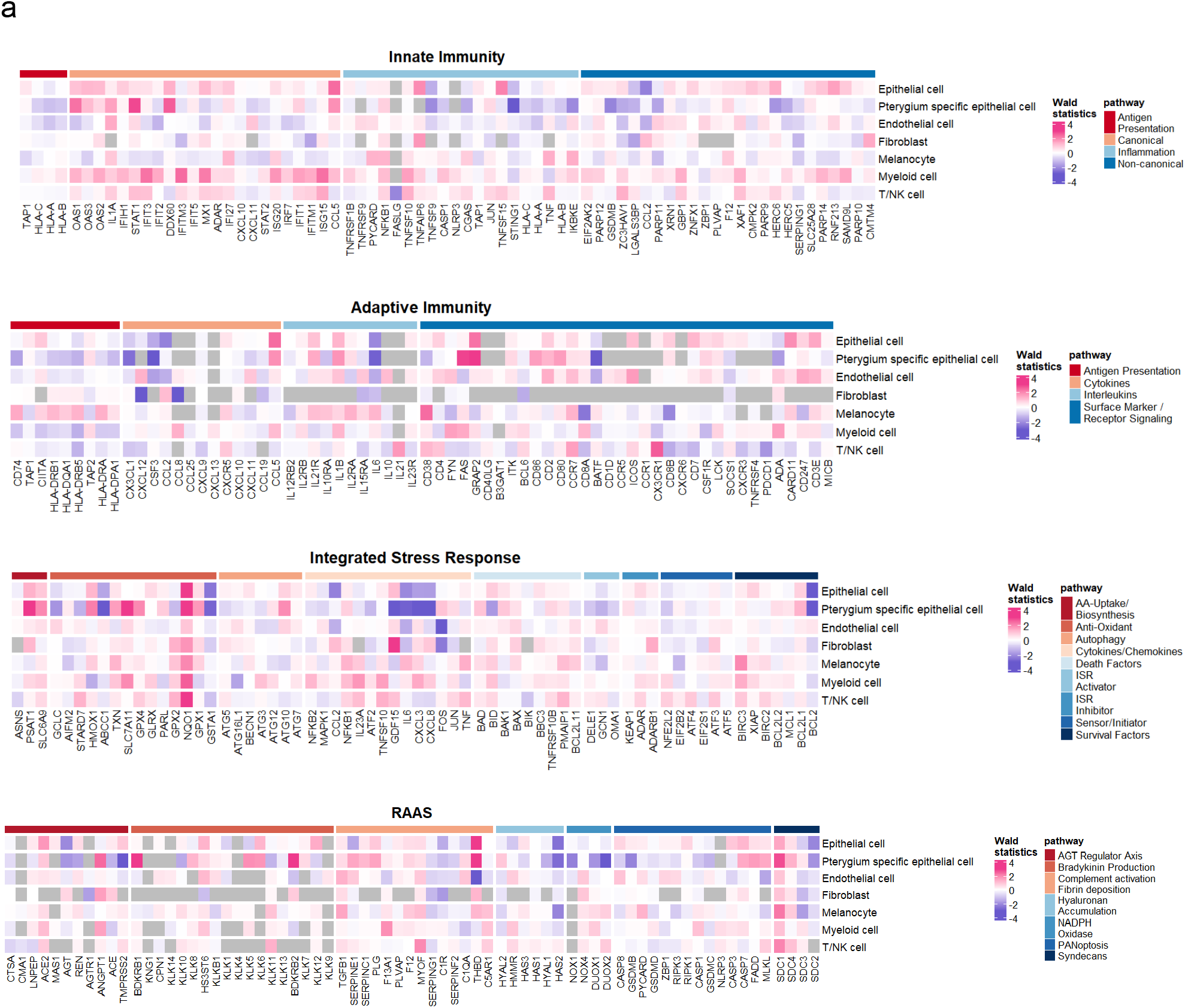
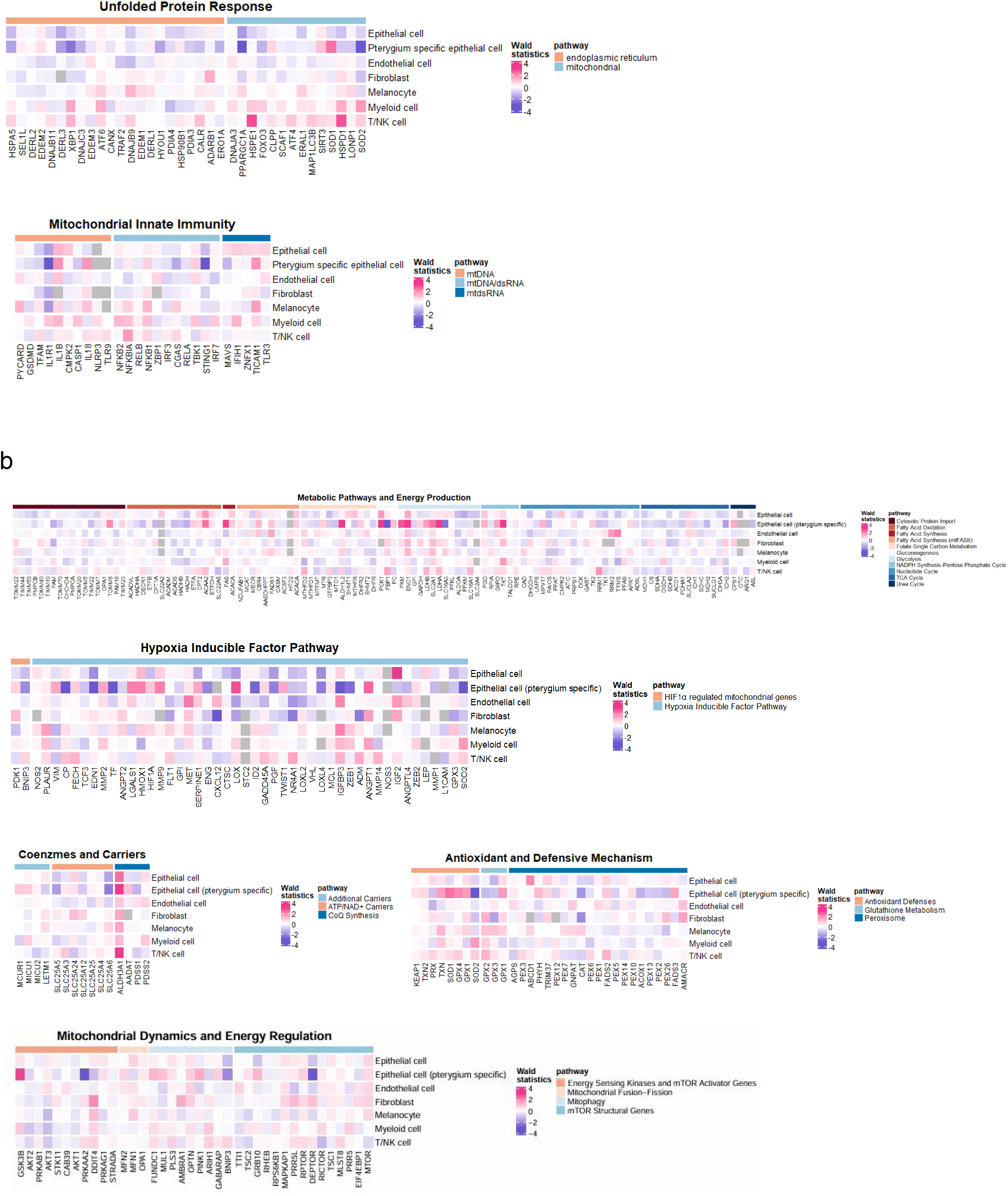
**(a)** Heatmaps of genes associated with inflammatory and **(b)** mitochondrial energy pathways. Wald statistics show upregulation (red) and downregulation (blue) of gene expression.

The significantly altered genes related to mitochondrial energy metabolism that were upregulated include: (i) PKM, ENO1, and LDHA, which reflect a shift toward aerobic glycolysis [21]; (ii) ALDH1L2 and TKT, which are involved in one-carbon metabolism and the pentose phosphate pathway [22]; (iii) SOD1, which mitigates oxidative damage in the cytoplasm. The significantly altered genes related to mitochondrial energy metabolism that were downregulated include: (i) FBP1 and SLC16A3, which are a gluconeogenic enzyme and a lactate transporter, respectively, suggesting an increased reliance of cells on glycolysis [23, 24]; (ii) SOD2, which is a mitochondrial superoxide dismutase, further confirming mitochondrial dysfunction [25] (Fig. 6b). These results align with the lollipop plot (Fig. 5) which showed metabolic reprogramming towards metabolic pathways.

Other cell types also contribute to the pathogenesis of pterygium. For instance, GDF15, known to modulate inflammation and promote tissue repair, is upregulated in fibroblasts, aligning with the fibrotic characteristics of pterygium. However, the downregulation of CXCL12 and CCL8 in fibroblasts suggests a reduced capacity for immune cell recruitment, potentially perpetuating unresolved inflammation. As CXCL12, along with its receptor CXCR4, regulates cell trafficking and immune responses, its downregulation could contribute to impaired tissue repair and persistent inflammation [26].

In T/NK cells, the upregulation of oxidative stress response genes, including NQO1, HSPE1, and HSPD1, reflects the immune cells’ attempts to counteract oxidative damage in the pterygium microenvironment. The HSPE1 and HSPD1 proteins form a mitochondrial chaperonin system that prevents protein misfolding and maintains mitochondrial proteostasis. The upregulation of these genes thus suggests that immune cells are under significant oxidative stress, leading to a reliance on these proteins to maintain mitochondrial integrity and mitigate the effects of chronic stress [27].

## 3. DISCUSSION

The use of multi-omics approaches powered by computational analysis is becoming increasingly vital to address diverse biomedical research questions. While such an approach is highly valuable, selecting the appropriate pathological region for clinical research is essential. In pterygium, the Main region (head, and neck) is involved in multiple inflammatory reactions that damage the cornea and is the central focus of surgical excision [28]. We began our analysis by separating the Main region from the Acc (body) region of pterygium and identified significant transcriptional profile differences between the two regions. A comprehensive PCA showed a clear separation of Acc from the Main and the Normal groups. Further analysis using databases such as GO and GSEA, along with custom inflammatory gene lists and Mitocarta 3.0, confirmed the distinct transcriptomic profiles between the Main and Acc regions. This distinction suggests the need for a greater emphasis on the anatomical regions of pterygium in future studies.

To further investigate the similar transcriptomic profiles between the Main and Normal groups, we utilized dissociation-based single-cell omics technologies that characterize cellular identities and tissue states, enabling us to examine the molecular and cellular landscape in patient-matched selective pterygium samples. Due to the transcriptional differences we identified within the anatomical regions of pterygium, we selectively collected the Main samples (non-ACC) to represent pterygium. Clustering of cells according to the cell types revealed pterygium-specific epithelial cells, a group of epithelial cells that were highly involved in the pathology of pterygium. Using CellChat, we observed that these epithelial cells interacted with fibroblasts, myeloid cells, endothelial cells, and other epithelial cells through the ANGPTL, IL-1, and KLK signaling pathways. The ANGPTL pathway, involving angiopoietin-like proteins (ANGPTLs), is known to regulate angiogenesis, cell migration, and inflammation [29]. Specifically, ANGPTL4, expressed in the conjunctival epithelium of surgically excised pterygia, is known to act as a secondary HIF-regulated angiogenic mediator [30]. The IL-1 signaling pathway, essential for immune responses, regulates inflammation, immune activity, and host defense, with studies showing overexpression of IL-1 in pterygium compared to normal conjunctiva [31]. The KLK signaling network, regulated by kallikrein-related peptidases (KLKs), involves proteolysis, inflammation, wound healing, and tissue remodeling. KLKs activate proteases, growth factors, and cytokines, playing critical roles in diseases such as cancer and skin disorders [31]. This suggests that the pterygium-specific epithelial cells found in this study are highly related to the immune alterations of pterygium identified in prior studies.

To reveal the sources of energy that the inflammatory pathways upregulated in pterygium receive, we analyzed the correlation between the inflammatory and mitochondrial energy pathways. Two inflammatory pathways, AA-uptake/biosynthesis in ISR and bradykinin production in RAAS, showed the highest correlation with the energy pathways, suggesting that the two use a great amount of energy in pterygium. A further analysis of the pathways specific to the cell types revealed that the expression of these two pathways was mainly increased in the pterygium-specific epithelial cells, matching our hypothesis that the cell group plays a major role in the expression of pterygium. The other mitochondrial energy pathways that showed significant correlations with the inflammatory pathways were metabolic pathways, MCD, and OXPHOS. When we further looked into the expression of energy metabolism-related genes using Mitocarta 3.0, pterygium samples were downregulated in the OXPHOS and MCD pathways compared to the control samples. The metabolic pathways, on the other hand, were upregulated, suggesting a metabolic reprogramming towards metabolic pathways in pterygium. This pattern corresponds to a previous study that reported the downregulation of OXPHOS in Asian samples [2].

Further confirming the role of pterygium-specific epithelial cells, we found that most of the significantly altered genes were present within the cell group in our analysis of the expression of specific genes. The pterygium-specific epithelial cells demonstrated an upregulation of key pro-inflammatory and stress-related genes. Specifically, STAT1 and DDX60, both crucial for type I interferon signaling, were significantly elevated, suggesting a sustained pro-inflammatory response. STAT1 plays a critical role in immune responses, particularly in antiviral defense, where its activation by interferons leads to the transcription of genes involved in immune signaling, apoptosis, and stress responses [12, 13]. This contributes to a pro-inflammatory environment and potential immune activation, reflecting chronic exposure to stressors like UV radiation. Additionally, the elevated expression of FAS, a key marker that triggers apoptosis through interactions with the FAS ligand (FASL), suggests increased cell death and tissue remodeling in pterygium. The active removal of damaged cells, likely driven by chronic inflammation, contributes to the degeneration and regrowth of epithelial layers [14].

Moreover, the increased expression of SLC7A11 and NQO1, both involved in antioxidant defense mechanisms, suggests that pterygium-specific epithelial cells are experiencing oxidative stress, likely triggered by chronic UV exposure. SLC7A11 is the functional light chain subunit of the cystine/glutamate antiporter system x_c_ ^−^, responsible for cystine uptake, which is critical for glutathione biosynthesis. Given its role in maintaining the cellular redox environment by promoting glutathione production, the upregulation of SLC7A11 indicates a heightened need for antioxidant defenses in response to chronic oxidative stress [15]. NQO1, heavily involved in redox processes, detoxifies reactive quinones and reduces reactive oxygen species (ROS), underscoring the oxidative stress response [16]. The upregulated expressions of these genes highlight the efforts to counter oxidative damage in pterygium cells, a key factor that drives both chronic inflammation and abnormal tissue remodeling in pterygium [32].

The increased expression of BDKRB1 and BDKRB2, which encode bradykinin receptors, suggests an inflammatory environment in pterygium caused by promoting vascular permeability, pain, and tissue swelling. Furthermore, the expression of THBD (thrombomodulin) and SDC1 (syndecan-1) points to ongoing tissue repair and remodeling. As SDC1 plays a critical role in wound healing, including cell adhesion, migration, and fibrosis, its upregulation underscores its involvement in the characteristic fibrotic processes in pterygium [17].

Conversely, the downregulation of CX3CL1, CSF3, IL6, and CXCL8 suggests an overall impairment in immune responses, potentially contributing to chronic inflammation. CX3CL1, for instance, is a transmembrane protein that binds to the CX3CR1 receptor, mediating leukocyte adhesion and the recruitment of immune cell subpopulations. Its downregulation reflects impaired immune cell recruitment and adhesion, which may perpetuate inflammation in pterygium tissues [18]. CSF3 (also known as G-CSF) promotes neutrophil survival and differentiation, leading to a diminished capacity for immune surveillance and tissue repair when reduced in expression [19]. This likely contributes to persistent inflammation and reduced tissue repair, creating a permissive environment for the progression of pterygium.

The downregulation of mitochondrial-related genes such as PPARGC1A (PGC-1α) and SOD2 further supports mitochondrial dysfunction. PGC-1α, a master regulator of mitochondrial biogenesis and oxidative phosphorylation, is critical for the maintenance of mitochondrial functions and the detoxification of ROS. Its downregulation thus exacerbates oxidative stress, potentially explaining the mitochondrial dysfunction and inflammatory responses found in pterygium-specific epithelial cells [20].

The limitation of this study is the lack of consideration of the metabolomic aspects of pterygium pathology. The incorporation of metabolomic data, particularly focusing on the metabolites we identified as significant, would enrich the transcriptomic and proteomic analyses performed in this study. Additionally, although we identified numerous biomarkers, we were unable to validate them experimentally. Further studies are necessary to verify the roles of these biomarkers.

## 4. CONCLUSION

In conclusion, the different expression patterns between the Main and the Acc groups highlight the need for an anatomical distinction in future studies. Furthermore, the cell-cell communication signaling networks and the upregulated inflammatory pathways showed that the epithelial cells specific to the Main group of pterygium play a major role in the inflammation of pterygium. The overexpression of pro-inflammatory and antioxidant genes and the downregulation of immune genes in the epithelial cells underscored the oxidative stress the cells received. Overall, our findings suggest potential directions to enhance our understanding of the pathogenesis of pterygium.

## 5. METHODS

### 5.1. Sample Collection and Preprocessing

Each of the Main, ACC, and Normal samples of the bulk RNA-seq data was collected from six primary pterygium patients by elective pterygium excision, approved by Kyung Hee University Hospital at Gangdong (IRB No. 2022-04-006). The patients, aged 61 to 72, included three males and three females. The head and the neck of pterygium were collected separately, each denoted as Main and ACC, respectively. The conjunctival tissue from each patient was collected as a control, denoted as Normal. FASTQC (v0.11.7) was used to check the quality of the data, and STAR (v2.7.3a) and HTSeq-Count (v0.12.4) were used to map the reads to the reference genome (GRCH38). The data was normalized using the rlog (regularized log) transformation from the DESeq2 package (v1.44.0). Single-cell RNA sequencing data was collected by Kyung Hee University Hospital at Gangdong using the Chromium Next GEM Single Cell 3’ RNA library kit (v3.1). Both the Main and Normal samples were collected from each of the two pterygium patients in the dataset. FASTQC (v0.11.7) was performed on the raw sequences for a quality check. The FASTQ files were preprocessed through 10x Genomics Cloud Analysis by choosing Next GEM Single Cell 3’ Gene Expression as the library type and setting checkLibraryCompatibility to true, Chemistry to auto, createBam to false, disableAbAggregateDetection to false, includeIntrons to true, and noSecondaryAnalysis to false [33]. The resulting gene-barcode matrix output was downloaded as a Market Exchange (MEX) file format.

The Seurat R package (v5.1.0) was used to control the quality of the data and perform integration. The data was filtered to those with less than 9000 genes detected per cell and less than 10,000 unique molecular identifiers (UMIs) per cell. The data was normalized with the NormalizeData function using the LogNormalize method, and anchor-based canonical correlation analysis (CCA) integration was applied through the IntegrateLayers function.

### 5.2. Cell Type Annotation

The Seurat package (v5.1.0) was used to generate clusters of the cells. FindNeighbors and FindClusters were performed with a resolution of 0.6 and the first 12 principal components (PCs) were chosen based on the results of the elbow plot and the heatmap of the top 20 PCs. Marker genes from Zhang et al. were used along with the cell atlas of the human ocular anterior segment from the Broad Institute to annotate the clusters [10].

### 5.3. Differential and Enrichment Analysis

The differential gene expression within the bulk RNA-seq data was analyzed by performing PCA and EnhancedVolcano using the DESeq2 package (v1.44.0). A p-value cutoff of 0.05 and an FC cutoff of 0.38 were used in the volcano plots. The enrichGO function from the clusterProfiler R package (v4.13.3) was performed to search for enriched functional pathways. The top 10 common pathways were identified from each comparison group (Main vs ACC, Normal vs ACC), resulting in 16 unique pathways when merged as a bubble plot. PPI was also depicted by STRING using K-means clustering with the top 1000 differentially expressed genes. Gene Ontology was used to identify the biological processes related to cluster 1 from the PPI of both comparison groups.

For both the bulk and single-cell RNA-seq data, differential gene expressions were measured using the DESeq2 R package (v1.44.0) with the results filtered to those with p-values less than 0.5. For the analysis of inflammatory pathways, a list of genes from the Molecular Signatures Database (MSigDB) was applied in fGSEA to obtain the NES for each pathway. The list was obtained by searching for homo sapiens genes relevant to a biological process (BP) with the keyword “inflammation”. MitoCarta3.0 was utilized for the enrichment analysis of energy-related pathways. The mitochondrial gene list obtained from Guarnieri et al. was used for the gene expression profile heatmaps of the energy pathways [7].

### 5.4. Correlation Analysis

The correlation between the inflammatory and energy pathways was analyzed by the Van Der Waerden test, comparing the NES obtained from fGSEA of the pathways. The results from fGSEA were filtered to those with p-values less than 0.3, and the results from the Van Der Waerden test were filtered to those with p-values greater than 0.9 and number of observations greater than 3.

### 5.5. Cell-cell Communication Analysis

The CellChat R package (v2.1.2) was used to compare the interactions between cell types in the pterygium and control samples. The interactions were limited to those related to inflammation found in CellChatDB. ComputeCommunProb was used with the type as truncatedMean and the trim as 0.05. Communications with less than 10 cells in each cell group were filtered out using filterCommunication.

## Author Contributions

Conceptualization, M.S.K. (Man S. Kim) and M.S.K. (Min Seok Kang);

Investigation, M.S.K. (Man S. Kim) and M.S.K. (Min Seok Kang);

Formal Analysis, C.S., S.M. and H.C.;

Methodology, C.S., S.M. and H.C.;

Software, Y.-A.K. and J.K.;

Visualization, M.S., and H.Y.;

Resources, T.G.K.;

Supervision, M.S.K. (Man S. Kim);

Writing—original draft preparation, All authors;

Writing—review and editing, M.S.K. (Man S. Kim) and M.S.K. (Min Seok Kang);

All authors have read and agreed to the published version of the manuscript.

## Notes

### Competing Interest Statement

The authors have declared no competing interest.

## References

[1] Džunić, B., Jovanović, P., Veselinović, D., Petrović, A., Stefanović, I., & Kovačević, I. (2010). Analysis of pathohistological characteristics of pterygium. Bosnian Journal of Basic Medical Sciences, 10(4), 307–313. 10.17305/bjbms.2010.2677

[2] Kim, Y.-A., Choi, Y., Kim, T. G., Jeong, J., Yu, S., Kim, T., Sheen, K., Lee, Y., Choi, T., Park, Y. H., Kang, M. S., & Kim, M. S. (2024). Multi-system-level analysis with RNA-seq on pterygium inflammation discovers association between inflammatory responses, oxidative stress, and oxidative phosphorylation. International Journal of Molecular Sciences, 25(9), 4789. 10.3390/ijms25094789

[3] Ghiasian, L., Samavat, B., Hadi, Y., Arbab, M., & Abolfathzadeh, N. (2021). Recurrent pterygium. Journal of Current Ophthalmology, 33(4), 367–378. 10.4103/joco.joco_153_20

[4] Shahraki, T., Arabi, A., & Feizi, S. (2021). Pterygium: An update on pathophysiology, clinical features, and Management. Therapeutic Advances in Ophthalmology, 13. 10.1177/25158414211020152

[5] Akbari M. (2022). Update on overview of pterygium and its surgical management. Journal of Population Therapeutics and Clinical Pharmacology, 29(4). 10.47750/jptcp.2022.968

[6] Wolf, J., Hajdu, R. I., Boneva, S., Schlecht, A., Lapp, T., Wacker, K., Agostini, H., Reinhard, T., Auw-Hädrich, C., Schlunck, G., & Lange, C. (2022). Characterization of the cellular microenvironment and novel specific biomarkers in Pterygia using RNA sequencing. Frontiers in Medicine, 8. 10.3389/fmed.2021.714458

[7] Guarnieri, J. W., Topper, M. J., Beigel, K., Haltom, J. A., Chadburn, A., Frere, J., An, J., Cope, H., Borczuk, A., Sinha, S., Lim, C., Kim, J., Park, J., Meydan, C., Foox, J., Mozsary, C., Bram, Y., Richard, S., Epsi, N. J., … Baylin, S. B. (2023). Lethal Covid-19 Associates with Raas-Induced Inflammation for Multiple Organ Damage Including Mediastinal Lymph Nodes. 10.1101/2023.10.08.561395

[8] Patergnani, S., Bouhamida, E., Leo, S., Pinton, P., & Rimessi, A. (2021). Mitochondrial oxidative stress and “mito-inflammation”: Actors in the diseases. Biomedicines, 9(2), 216. 10.3390/biomedicines9020216

[9] Ransy, C., Vaz, C., Lombès, A., & Bouillaud, F. (2020). Use of H2O2 to cause oxidative stress, the catalase issue. International Journal of Molecular Sciences, 21(23), 9149. 10.3390/ijms21239149

[10] Zhang, X., Han, P., Qiu, J., Huang, F., Luo, Q., Cheng, J., Shan, K., Yang, Y., & Zhang, C. (2024). Single-cell RNA sequencing reveals the complex cellular niche of pterygium. The Ocular Surface, 32, 91–103. 10.1016/j.jtos.2024.01.013

[11] Wilkinson-Berka, J. L., Suphapimol, V., Jerome, J. R., Deliyanti, D., & Allingham, M. J. (2019). Angiotensin II and aldosterone in retinal vasculopathy and inflammation. Experimental Eye Research, 187, 107766. 10.1016/j.exer.2019.107766

[12] Tolomeo, M., Cavalli, A., & Cascio, A. (2022). STAT1 and its crucial role in the control of viral infections. International Journal of Molecular Sciences, 23(8), 4095. 10.3390/ijms23084095

[13] Miyashita, M., Oshiumi, H., Matsumoto, M., & Seya, T. (2011). DDX60, a DEXD/H box helicase, is a novel antiviral factor promoting rig-i-like receptor-mediated signaling. Molecular and Cellular Biology, 31(18), 3802–3819. 10.1128/mcb.01368-10

[14] Strasser, A., Jost, P. J., & Nagata, S. (2009). The many roles of Fas receptor signaling in the immune system. Immunity, 30(2), 180–192. 10.1016/j.immuni.2009.01.001

[15] Hu, K., Li, K., Lv, J., Feng, J., Chen, J., Wu, H., Cheng, F., Jiang, W., Wang, J., Pei, H., Chiao, P. J., Cai, Z., Chen, Y., Liu, M., & Pang, X. (2020). Suppression of the SLC7A11/glutathione axis causes synthetic lethality in kras-mutant lung adenocarcinoma. Journal of Clinical Investigation, 130(4), 1752–1766. 10.1172/jci124049

[16] Ross, D., & Siegel, D. (2021). The diverse functionality of NQO1 and its roles in redox control. Redox Biology, 41, 101950. 10.1016/j.redox.2021.101950

[17] Stepp, M. A., Pal-Ghosh, S., Tadvalkar, G., & Pajoohesh-Ganji, A. (2015). Syndecan-1 and its expanding list of contacts. Advances in Wound Care, 4(4), 235–249. 10.1089/wound.2014.0555

[18] Rivas-Fuentes, S., Salgado-Aguayo, A., Arratia-Quijada, J., & Gorocica-Rosete, P. (2021). Regulation and biological functions of the CX3CL1-CX3CR1 axis and its relevance in solid cancer: A mini-review. Journal of Cancer, 12(2), 571–583. 10.7150/jca.47022

[19] Garg, B., Mehta, H. M., Wang, B., Kamel, R., Horwitz, M. S., & Corey, S. J. (2020). Inducible expression of a disease-associated ELANE mutation impairs granulocytic differentiation, without eliciting an unfolded protein response. Journal of Biological Chemistry, 295(21), 7492–7500. 10.1074/jbc.ra120.012366

[20] Rius-Pérez, S., Torres-Cuevas, I., Millán, I., Ortega, Á.L., & Pérez, S. (2020). PGC-1a, inflammation, and oxidative stress: An integrative view in metabolism. Oxidative Medicine and Cellular Longevity, 2020, 1–20. 10.1155/2020/1452696

[21] Zhang, Z., Deng, X., Liu, Y., Liu, Y., Sun, L., & Chen, F. (2019). PKM2, function and expression and regulation. Cell & Bioscience, 9(1). 10.1186/s13578-019-0317-8

[22] Kim, Y., Kim, E.-Y., Seo, Y.-M., Yoon, T. K., Lee, W.-S., & Lee, K.-A. (2012). Function of the pentose phosphate pathway and its key enzyme, transketolase, in the regulation of the meiotic cell cycle in oocytes. Clinical and Experimental Reproductive Medicine, 39(2), 58. 10.5653/cerm.2012.39.2.58

[23] Park, H.-J., Jang, H. R., Park, S.-Y., Kim, Y.-B., Lee, H.-Y., & Choi, C. S. (2020). The essential role of fructose-1,6-bisphosphatase 2 enzyme in thermal homeostasis upon cold stress. Experimental & Molecular Medicine, 52(3), 485–496. 10.1038/s12276-020-0402-4

[24] Tao, Q., Li, X., Zhu, T., Ge, X., Gong, S., Guo, J., & Ma, R. (2022). Lactate transporter SLC16A3 (MCT4) as an onco-immunological biomarker associating tumor microenvironment and immune responses in lung cancer. International Journal of General Medicine, Volume 15, 4465–4474. 10.2147/ijgm.s353592

[25] Flynn, J. M., & Melov, S. (2013). SOD2 in mitochondrial dysfunction and neurodegeneration. Free Radical Biology and Medicine, 62, 4–12. 10.1016/j.freeradbiomed.2013.05.027

[26] Mezzapelle, R., Leo, M., Caprioglio, F., Colley, L. S., Lamarca, A., Sabatino, L., Colantuoni, V., Crippa, M. P., & Bianchi, M. E. (2022). CXCR4/CXCL12 activities in the tumor microenvironment and implications for tumor immunotherapy. Cancers, 14(9), 2314. 10.3390/cancers14092314

[27] Yeung, N., Murata, D., Iijima, M., & Sesaki, H. (2023). Role of human HSPE1 for OPA1 processing independent of HSPD1. iScience, 26(2), 106067. 10.1016/j.isci.2023.106067

[28] Khandelwal, R., Tigga, M., Metri, R., & Deshpande, A. (2024). Comparative study of pterygium excision with suture and sutureless conjunctival autograft. European Journal of Clinical and Experimental Medicine, 22(2), 334–339. 10.15584/ejcem.2024.2.15

[29] Carbone, C., Piro, G., Merz, V., Simionato, F., Santoro, R., Zecchetto, C., Tortora, G., & Melisi, D. (2018). Angiopoietin-like proteins in angiogenesis, inflammation and cancer. International Journal of Molecular Sciences, 19(2), 431. 10.3390/ijms19020431

[30] Meng, Q., Qin, Y., Deshpande, M., Kashiwabuchi, F., Rodrigues, M., Lu, Q., Ren, H., Elisseeff, J. H., Semenza, G. L., Montaner, S. V., & Sodhi, A. (2017). Hypoxia-Inducible Factor-Dependent Expression of Angiopoietin-Like 4 by Conjunctival Epithelial Cells Promotes the Angiogenic Phenotype of Pterygia. Invest Ophthalmol Vis Sci, 58(11), 4514–4523. 10.1167/iovs.17-21974

[31] Zhou, W. P., Zhu, Y. F., Zhang, B., Qiu, W. Y., & Yao, Y. F. (2016). The role of ultraviolet radiation in the pathogenesis of pterygia (Review). Mol Med Rep, 14(1), 3–15. 10.3892/mmr.2016.5223

[32] Suarez, M. F., Echenique, J., López, J. M., Medina, E., Irós, M., Serra, H. M., & Fini, M. E. (2021). Transcriptome analysis of pterygium and Pinguecula reveals evidence of genomic instability associated with chronic inflammation. International Journal of Molecular Sciences, 22(21), 12090. 10.3390/ijms222112090

[33] Cloud analysis. 10x Genomics. https://www.10xgenomics.com/support/software/cloud-analysis/latest

